# Phosphorylation of Ube2J1 at serine S184 is regulated by protein phosphatase 2A

**DOI:** 10.64898/2026.03.30.715004

**Authors:** Dominic S. Dollken, Shuet Y. Lam, Tymoteusz K. Kaminski, John V. Fleming

**Author notes:** Correspondence: Eoin (J.V.) Fleming, School of Biochemistry and Cell Biology and School of Pharmacy, University College Cork, Cork, Ireland.

## Abstract

The Ube2J1 enzyme that mediates the ubiquitination and proteasomal degradation of misfolded proteins at the ER is phosphorylated at serine S184. Following anisomycin treatment of HEK293T cells, we observed an inverse relationship between phosphorylation and dephosphorylation at this site. This suggested a dynamic interchange between the two forms, and we show that S184 is a target for protein phosphatase 2A. The S184-phosphorylated protein is known to exhibit increased sensitivity to proteasomal degradation, and we found that mutation at K186R increased the ratio of S184-phosphorylated to S184-dephosphorylated protein. Although the K186R mutant retained some sensitivity to proteasomal inhibition, our results show that Ube2J1 steady state expression can be exercised at multiple levels, and can involve dynamic phosphorylation and dephosphorylation at S184.

## INTRODUCTION

Protein ubiquitination is an important post-translational modification that regulates numerous cellular processes. It is achieved catalytically by the sequential action of an E1 ubiquitin-activating enzyme, an E2 ubiquitin-conjugating enzyme, and an E3 ubiquitin ligase (1). It is estimated that the human genome encodes two E1s, approximately 40 E2s, and >600 different E3s (2-4). Lysine is considered the canonical site for covalent attachment of ubiquitin to substrates, with attachment occurring at the ε-amino group of the side chain. However, cysteine, serine and threonine residues, as well as the free amino group of the N-terminus of proteins, also function as sites for ubiquitination (5, 6).

In eukaryotic cells, proteostasis involves the folding and refolding of proteins at the endoplasmic reticulum to achieve the correct functional structure. Genetic and environmental conditions can influence and compromise this process and give rise to cellular conditions of persistent protein misfolding (7). In these cases, an evolutionarily conserved stress response referred to as the Unfolded Protein Response (UPR) is induced. This aims to minimize proteotoxicity, and promotes ER-associated degradation (ERAD) to ubiquitinate and proteasomally degrade misfolding protein variants (8). ER localized chaperones and lectins bind misfolding proteins and facilitate their retro-translocation through an ER membrane complex referred to as a translocon. E2 and E3 enzymes that are localized or recruited to the ER then ubiquitinate the misfolded variant (9, 10).

In mammalian cells, the ER-localized Ube2J1 (Ubc6e) enzyme plays a central role in ERAD, functioning alongside HRD1 (E3-ligase), SEL1L and Herp1 in one of the best described translocons (11-14). In this context it is known to contribute towards the degradation of several misfolding protein variants (11). In response to pharmacological induction of the UPR, the protein undergoes phosphorylation at serine residue S184 (14). Although additional studies will be required to fully characterize the effect of this on substrate ubiquitination, regulation at the site is reported to influence the affinity of the enzyme for the cIAP1 E3 ligase (15).

Although Ube2J1 functions in the degradation of substrate proteins, little is known about how the protein itself is degraded. Exchanging the carboxy-terminal tail that mediates ER localization for the corresponding region of other tail anchored proteins, has been shown to confer novel degradation properties, including ubiquitination and proteasomal degradation (16). While this suggests that the region is important for regulating Ube2J1 protein stability, the processes that normally regulate its degradation remain unclear, however it is evident that phosphorylation at S184 creates a less stable form of the protein that leads to increased degradation by the proteasome (15).

Here we identified lysine residue K186 as being of importance for regulating the steady state expression of the S184-phosphorylated form of the protein, and show that accumulation of the S184 form of the protein occurs as a result of multiple cellular processes, including phosphorylation, de-phosphorylation involving PP2A, and proteasomal degradation.

## MATERIALS AND METHODS

### Plasmid DNA

The coding sequence for human Ube2J1 was PCR amplified from a human testes cDNA library and cloned into the pEP7-HA and pEP7-His vectors as previously described (17, 18). The coding sequence for human HRD1/ Synoviolin and the α-catalytic subunit of human PP2A was generously gifted (19, 20) and PCR amplified for cloning into the pEP7-FLAG vector (21). PP2A was also cloned into the pTetOne-FLAG vector. The QuickChange protocol (Stratagene) was used to mutate the Ube2J1 coding sequence.

### Cell culture and transfections

HEK293T (ATCC) were seeded at densities of 1 × 10^5^ cells per well on 24 well plates and transfected with Lipofectamine 2000 (Invitrogen) as described elsewhere (21). Cells were incubated in the presence or absence of pharmacological agents purchased from Sigma (MG132, okadaic acid, doxycycline) as described in figure legends. Cells were harvested in RIPA buffer as described elsewhere (22).

### Immunoblot analysis

Lysates were fractionated on denaturing 11% SDS–polyacrylamide gels for immunoblotting by standard methods with mouse anti-FLAG (Sigma), anti-HA (Covence) and bespoke rabbit anti-Ube2J1 or rabbit anti-phospho Ube2J1-S184P (1:1000) antibodies (Davids Biotechnologie). Equal loading of gels was confirmed using mouse anti-α-tubulin or mouse anti-β-actin antibodies (Sigma). All immunoblots shown are representative of at least 3 independent experiments. Immunoblot imaging and densitometric analysis were performed on an iBright FL1500 imaging system (Thermo Fisher Scientific). Statistical analysis was performed in Graphpad using a Students paired t-test or one-way Anova with Dunnett’s post hoc test.

## RESULTS

### Anisomycin treatment differentially regulates higher and lower molecular weight forms of Ube2J1

Phosphorylation of the Ube2J1 protein at serine S184 can occur in response to several different stimuli, including p38 MAP kinase signaling (12). To better explore the dynamics of this phosphorylation, HEK293T cells were transfected to express Ube2J1 and treated pharmacologically with anisomycin, which acts primarily through p38 to regulate intracellular signalling events. As can be seen in Fig.1A, when cell lysates were analysed using an anti-Ube2J1 specific antibody we detected major higher and lower molecular weight bands (arrowheads). As can be further demonstrated in Fig.1A, anisomycin treatment resulted initially in a decrease in steady state levels of the lower band. This was matched with a corresponding increase in the higher molecular weight form. Between 1 hr and 3 hr post-stimulation this process was reversed, with decreases in the higher molecular weight form being mirrored by increases in the lower band. These dynamic changes were confirmed in changes to the ratio of the two major bands (Fig.1B).

**Fig. 1.**
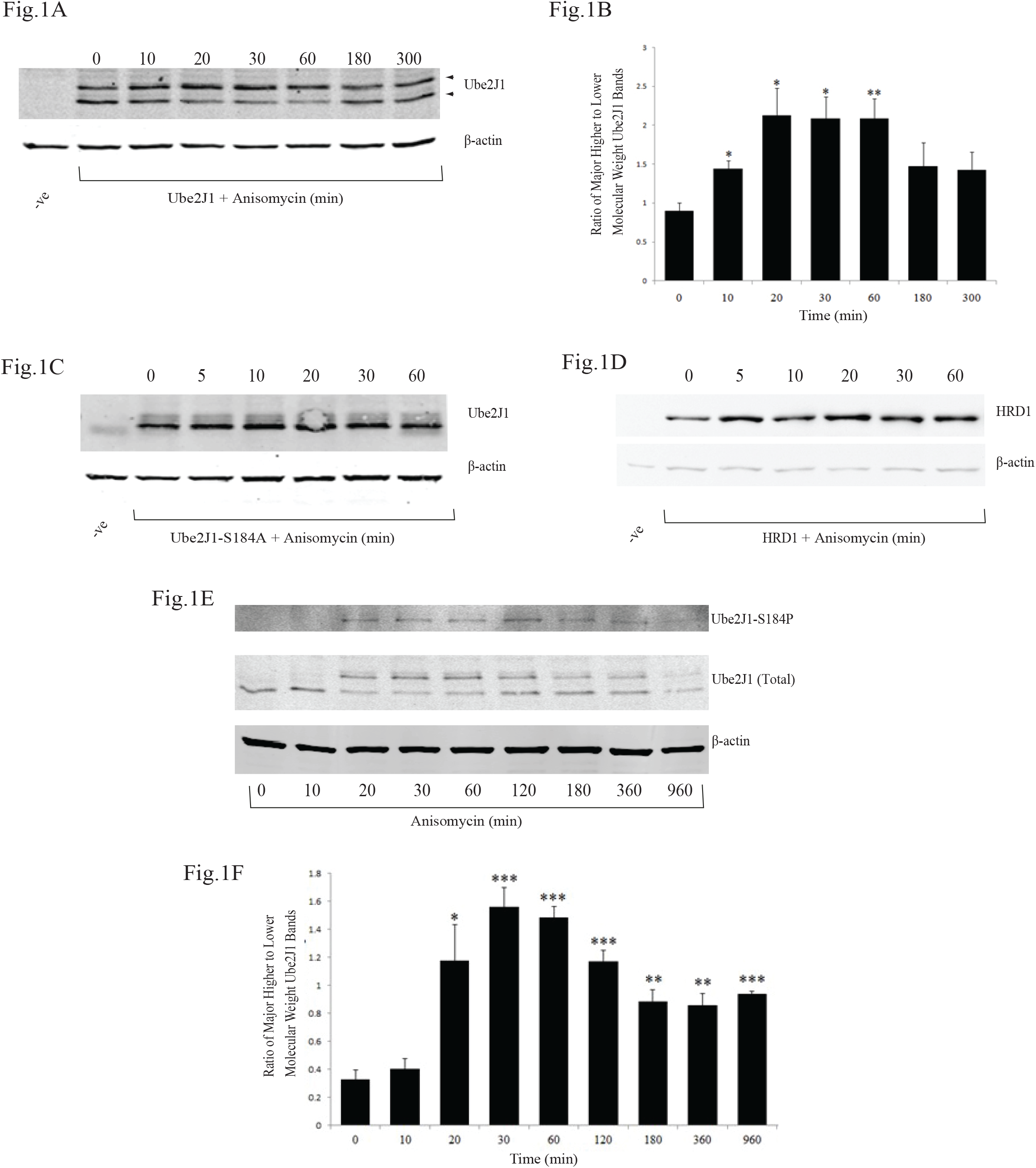
Ube2J1 is regulated by anisomycin. (A) HEK293T cells were transiently transfected to express Ube2J1 HA and serum starved overnight (16 h) before treating with anisomycin (50ng/ml) for the indicated times (minutes). Cells were harvested 48 h post-transfection and lysates were analysed on immunoblots using an anti-Ube2J1 antibody. (B) Bar graph showing densitometric quantification of the higher molecular weight band divided by the lower molecular weight band of Ube2J1 (arrowheads in Fig. 1A). Error bars are the SEM of three independent experiments. (C) HEK293T cells were transiently transfected to express Ube2J1-S184A HA before serum starvation and treatment with anisomycin (50ng/ml) at the indicated (minutes). The lysates were analysed on immunoblots to detect Ube2J1 using an anti-HA antibody. (D) HEK293T cells transiently transfected to express HRDl FL were serum starved and treated with anisomycin (50ng/ml) (minutes). The lysates were analysed on immunoblots to detect HRDl using an anti-FLAG antibody. (E) HEK293T cells were serum starved overnight (16 h) before treating with anisomycin (50ng/ml) for the indicated times (minutes). The lysates were analysed on immunoblots using anti-phospho UbeJ1-S184P (upper panel) or anti Ube2J1 antibodies (middle panel). (F) Bar graph showing densitometric quantification of the S184 phosphorylated band divided by the S184 dephosphorylated bands of Ube2J1 as described in Fig. 1E. Error bars are the SEM of three independent experiments. β-actin was used for loading controls. The significance of changes in the S184-phosphorylated to S184-dephosphorylated ratios were assessed by student T-test following comparison with the untreated controls (0 min),* p≤0.05, ** p≤0.01, *** p≤0.001

The S184-phosphorylated form of Ube2J1 is known to be less stable (15). Assuming that the higher band on our gels was indicative of S184, our result could be partly explained on the basis that the phosphorylated form of the protein was being sent for degradation following anisomycin treatment. While this would explain the decrease we observed in the higher molecular weight band at later time points, it would not explain the latter stage increases observed in the lower band.

To get a better understanding of the inverse relationship between the two bands, we tested whether the higher molecular weight band did indeed represent the S184-phosphorylated protein. HEK293T cells were transfected to express a S184A mutant form of the protein, and the treatment with anisomycin was repeated. As can be seen in Fig1C, the mutation resulted in loss of the higher molecular weight band, confirming that it represented the S184 phosphorylation. In its absence, the protein was not responsive to anisomycin in the same way as the wild type protein, and we failed to see any decrease through the 20-to-60-minute time points. This suggested that the changes in the lower molecular weight band observed in Fig1.A (decrease followed by increase) were occurring as a result of the ability of the protein to become phosphorylated. The HRD1 E3 ligase that functions with Ube2J1 in the translocon is not known to undergo p38 mediated phosphorylation. As can be seen in Fig.1D, there was no evidence for a decrease in protein levels in response to anisomycin.

We wished to test the cellular relevance of our observations and establish whether the endogenous Ube2J1 protein might be similarly regulated following anisomycin treatment of untransfected HEK293T cells. Using an anti-phospho S184 (S184P) antibody that preferentially recognizes the phosphorylated form of the protein, there was an initial accumulation of the phospho band in response to the drug. This reduced at later time points (Fig.1E). Using the antibody that detects total Ube2J1 protein, we were able to demonstrate once again that this was happening in the context of an inverse correlation between the higher and lower molecular bands that was reversed following 3-5hrs (Fig.1F). It was evident from these combined data therefore that a similar pattern of expression was being observed for both endogenous and ectopically expressed protein.

### The S184 phosphorylated protein accumulates with inhibition of the proteasome and inhibition of protein phosphatases

Previous studies suggest that the Ube2J1 protein is comparatively unstable and degraded by the proteasome (15). In our HEK293T cells model, the treatment of transfected cells for up to 16hr with the proteasome inhibitor MG132 led to an increase in total Ube2J1 protein levels (Fig.2A). Although our studies suggested that the lower molecular weight band increased in response to this treatment, it was evident that the S184 phosphorylated band was comparatively much more sensitive, and at a 5 hr timepoint (Fig.2B), there was a significant increase in the ratio of the higher to the lower bands (Fig.2C).

**Fig. 2.**
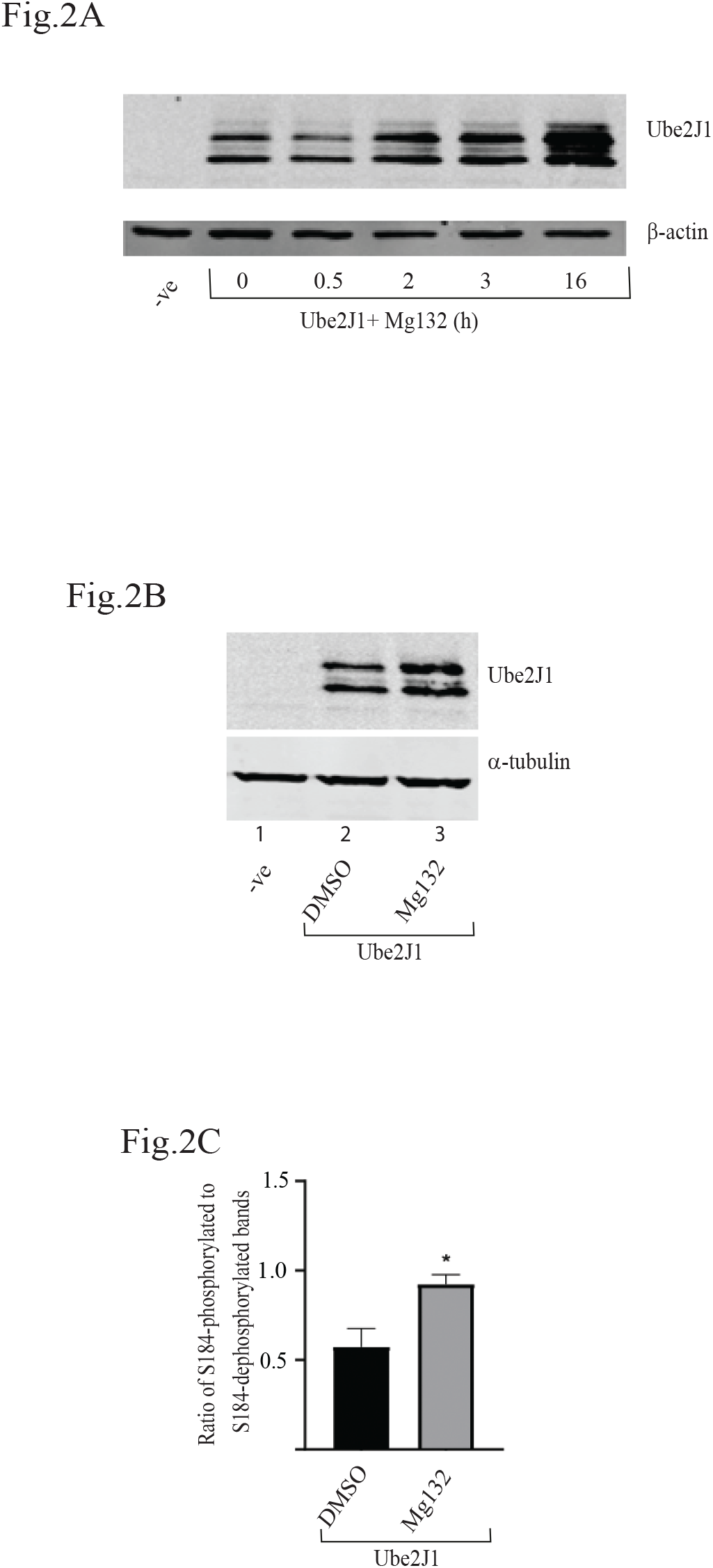
Ube2J1 is degraded by the proteasome. (A) HEK293T cells were transfected to transiently express Ube2J1-HA and treated for the indicated time periods in the presence or absence of MG132 at 20µM. -ve relates to cells transfected with empty plasmid vector (pcDNA 3.1). Cells were harvested 48 h post-trasnfection and lysates were analysed on immunoblots using an anti-Ube2J1 antibody. The expression levels of β actin were used as the loading control. (B) HEK293T cells transiently expressing Ube2J1-HA were treated for 5 h in the presence or absence of Mg132 (20µM) or DMSO. α-tubulin were used as the loading control (C) Bar graph showing densitometric quantification of the S184-phosphorylated Ube2J1 band divided by the dephosphorylated band for cells treated for 5 h with MG132 or DMSO. Error bars are the SEM of three independent experiments. * denotes significance with respect to control cells (DMSO treated). * P <0.05.

Although both the anisomycin and Mg132 treatments increased steady state levels of the S184-phosphorylated band, the underlying patterns were quite different. For the former, our data pointed towards a dynamic interconversion between the two bands, while for the proteasome inhibitor, there was no evidence for this. Although dephosphorylation of S184-phosphorylated protein has not previously been considered, we hypothesized that the observed pattern following anisomycin treatment could reflect a controlled return to the lower molecular weight form. To explore this possibility, HEK293T cells transfected to express Ube2J1 were incubated in the presence and absence of okadaic acid, which is a pharmacological inhibitor of multiple protein phosphatase enzymes. As can be seen in Fig.3, 20hr of this treatment resulted in a clear accumulation of the higher molecular weight band, and use of the anti-phospho S184P antibody confirmed that the protein is indeed subject to the activity of phosphatases.

**Fig. 3.**
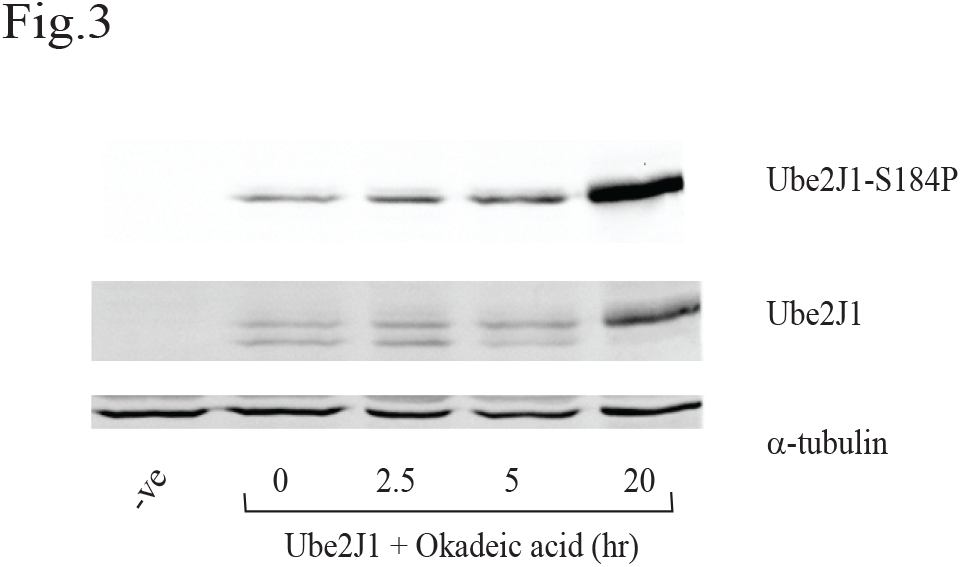
Treatment with okadaic acid leads to accumulation of the S184-phosphorylated form of Ube2J1. HEK293T cells were transfected to transiently express Ube2J1-His and treated for the indicated time periods in the presence or absence of okadaic acid (10nM). -ve relates to cells transfected with empty plasmid vector (pcDNA 3.1). Cells were harvested and lysed after 48 h of transfection and the lysates were analyzed on immunoblots using anti-phospho Ube2J1 S184P or anti Ube2J1 antibodies. α-Tubulin was used as the loading control.

### The S184 phosphorylation site of Ube2J1 is a target for PP2A

Okadaic acid acts to inhibit multiple phosphatases but acts preferentially on PP2A (23). To better understand this process, and help identify the specific phosphatase involved, HEK293T cells expressing Ube2J1 were co-transfected to express the α-catalytic subunit of PP2A (PP2A-α). As can be seen in Fig.4A this treatment led to loss of the higher molecular weight band, with our anti-phospho Ube2J1-S184P antibody confirming the loss of phosphorylation specifically at S184. In an independent model, we cloned PP2A-α downstream from a doxycycline regulated promoter. HEK293T cells were transfected to co-express a combination of Ube2J1 and PP2A-α expression constructs, and treated with increasing concentrations of doxycycline. As can be seen for Fig.4B, this led to regulated decreases in the S184 phosphorylated Ube2J1 band.

**Fig. 4.**
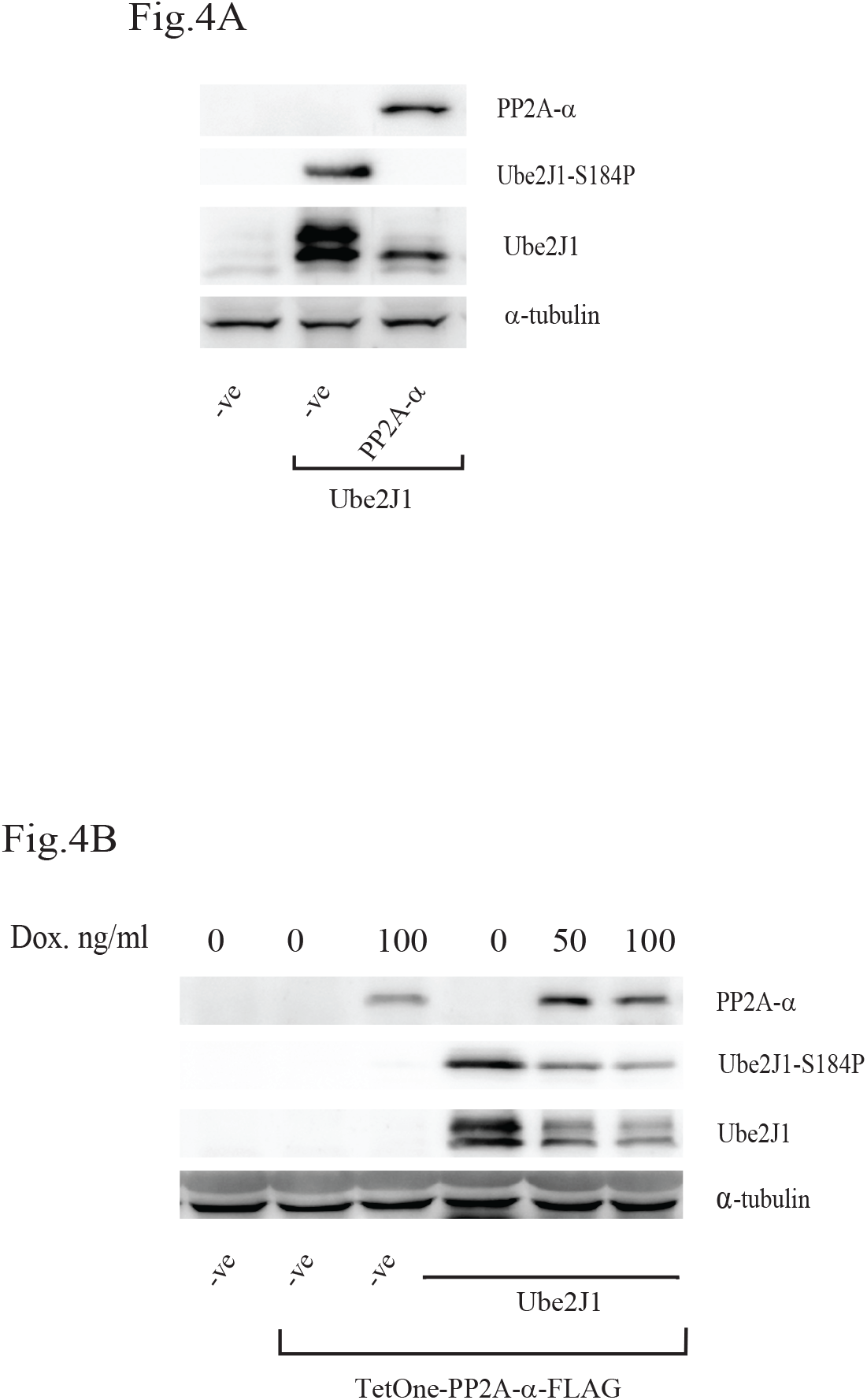
Co-expression of the α-catalytic subunit of PP2A decreases the accumulation of S184 phosphorylated Ube2J1. (A) HEK293T cells were transfected with pcDNA (-ve), Ube2J1-His, and PP2A-α-FLAG. Harvested lysates were analysed on immunoblots using anti-Ube2J1, anti-phospho Ube2J1-S184P, and anti-FLAG (to detect PP2A-α) antibodies. α-Tubulin was used as the loading control. (B) HEK293T cells were transfected with pcDNA (-ve), Ube2J1-His, and doxycycline inducible pTetOne-PP2A-α-FLAG for 72 h. Doxycycline was added to the indicated cells at the indicated concentrations for the 72 h of transfection. Cells were harvested and lysed after 72 h of transfection and the lysates were analysed on immunoblots using anti-Ube2J1, anti-Ube2J1-S184P and anti-FLAG (to detect PP2a-α) antibodies. α-Tubulin was used as the loading control.

### Lysine residue K186 is important for regulating steady state levels of the S184 phosphorylated form of Ube2J1

Together, these data suggest that steady state levels of S184-phosphorylated protein will reflect the dynamic phosphorylation, dephosphorylation and proteasomal degradation of Ube2J1. The amino acid lysine that is central to the process of proteasomal degradation, appears sixteen times in the Ube2J1 primary protein sequence. To determine whether any of these contribute towards the steady state expression of Ube2J1, we mutated them either individually or combinatorially (for residues found in close proximity to one another). The mutated constructs that were generated included: K8/13/17/23R, K71R, K88/89R, K122R, K142/143R, K157R, K164R, K177R, K186R, K194R, K246R. We also generated a Ube2J1 construct in which all of the lysine residues were mutated to arginine giving a “no lysine” (NoK) mutant.

To test whether lysine residues are important for regulating steady state expression of Ube2J1 isoforms, HEK293T cells were transfected to express wild type or mutant proteins. After harvesting and western blot analysis of lysates, we noted that the K186R and no lysine (NoK) mutants showed significant increases in the ratio of the S184-phosphorylated to dephosphorylated bands (Fig.5A lanes 11 and 14 respectively and Fig.5B).

**Fig. 5.**
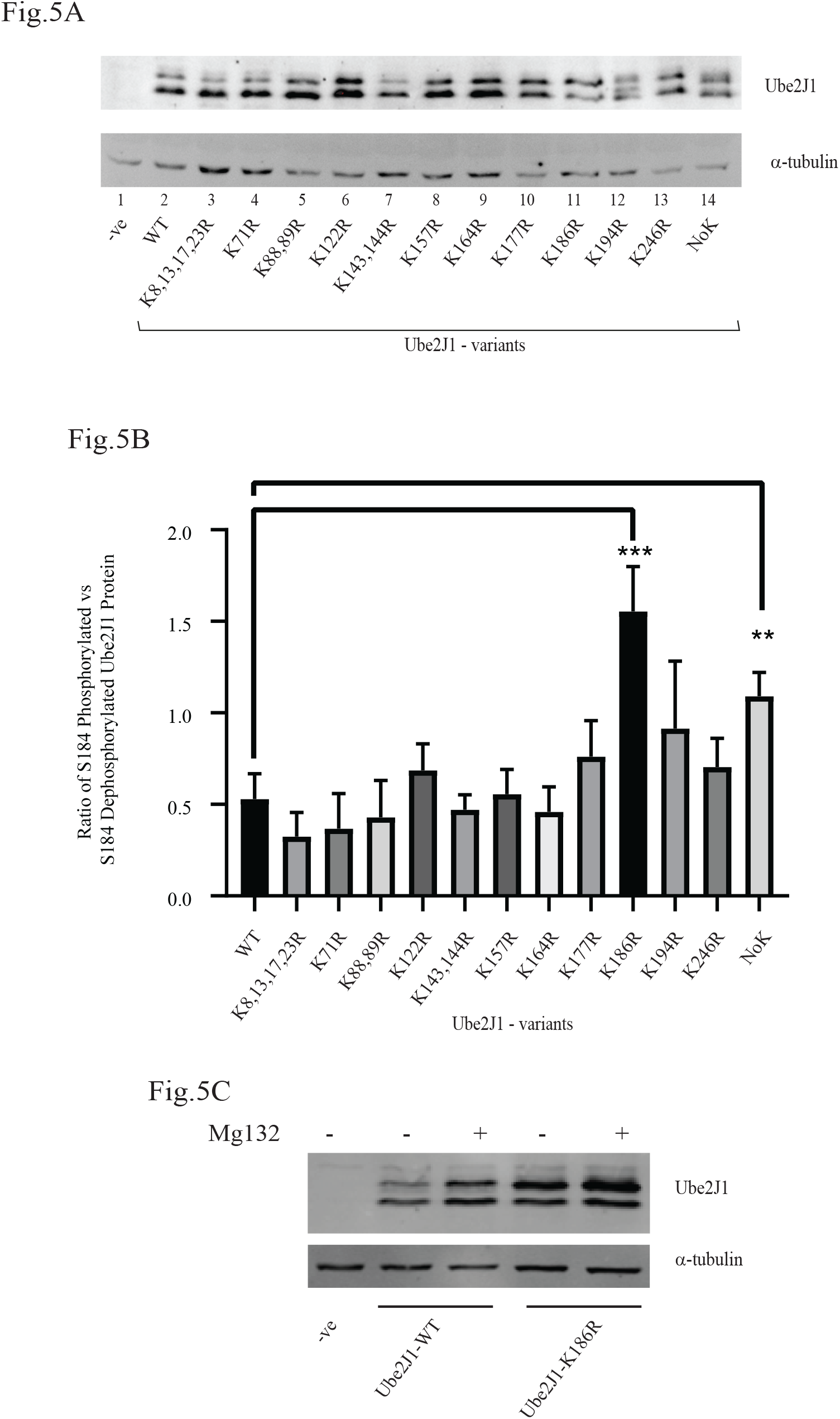
Lysine mutagenesis at K186 alters the ratio of S184-phosphorylated to S184-dephosphorylated Ube2J1. (a) HEK293T cells were transfected to transiently express empty vector (-ve), wild type Ube2J1-HA or Ube2J1-HA mutants with the indicated lysine residues replaced by arginines. The Ube2J1 -NoK-HA protein does not contain lysine residues. Cells were harvested 48 h after transfection and the lysates were analysed on immunoblots using an anti-Ube2J1 antibody. α-Tubulin was used as the loading control. (b) Bar chart representing the densitometric quantification of the S184-phosphorylated bands of Ube2J1 divided by the S184-dephosphorylated bands. Error bars are the SEM of three independent experiments. Statistical analysis was carried out comparing the band ratio between the wild type protein and each mutant. * denotes significance with respect to control cells (Ube2J1-WT-HA transfected). ** P<0.01, *** P< 0.001, calculated using one-way ANOVA with Dunnett’s test. (C) HEK293T cells were transfected to transiently express Ube2J1-HA variants as indicated and treated for 5 h in the presence or absence of MG132 at 20µM. -ve relates to cells transfected with empty plasmid vector. Cells were harvested after 48 h and lysates were analysed on immunoblots using an anti-Ube2J1 antibody. α-Tubulin was used as the loading control.

From our experiments presented in Fig.2, we were already conscious that the S184 phosphorylated form of Ube2J1 is less stable and more sensitive to proteasomal inhibition. Our observed changes in the ratio in Fig.5B could therefore have been due to a decrease in the degradation of the S184-phosphorylated protein, which resulted from the mutation at K186. In principle, this might be expected to make the mutant protein resistant to proteasomal degradation (and inhibition of the proteasome by the inhibitor MG132). To test this, cells transfected to express wild type or mutant protein were incubated in the presence and absence of Mg132. Following analysis, it was evident that the K186R mutant still responded to treatment with Mg132 (Fig.5C).

## DISCUSSION

Phosphorylation of the Ube2J1 is known to occur at serine S184. While this was first discovered in the context of the UPR, where it was shown to be downstream from PERK signalling (14), subsequent analysis has shown it to be a feature of a number of additional stress conditions (12). This pattern of regulation suggests that it functions to integrate several different signals, and its presence is important for a variety of cellular responses. In terms of S184 phosphorylation, our studies demonstrated that steady state expression will be a reflection of both phosphorylation and proteasomal degradation, however we also demonstrate a role for dephosphorylation, and the importance of a proximal K186 site. This points towards a broader regulation of the S184 phospho-protein, and the possibility for fine tuning of its steady state expression.

The significance of this on ERAD, and the emerging roles in which Ube2J1 has been additionally implicated (24-27), depends on how the S184 phosphorylation might influence enzyme function. It does not for example impact on cellular localization to the ER, and although there is evidence that S184D and S184E phospho-mimetics may have altered catalytic activity, it does not appear to influence interactions with the E3 ligase parkin, or impact on degradation of a target substrate (14). More recently, there have been reports for differential binding to the E3 ligase cIAP1, but once again it’s significance for the degradation of ERAD substrates is less clear. Instead, the clearest functional effect to this point relates to protein stability (15).

While our studies in HEK293T cells support previous reports (15) that S184-phosphorylation influences the stability of the protein by making it more susceptible to proteasomal degradation, we found little evidence to suggest that the sole purpose of phosphorylation is to promote its degradation. If this was the case, we might have expected for anisomycin to decrease total Ube2J1 protein levels over time. Instead, we observed a dynamic change in the ratio of the higher and lower molecular weight bands, which was reversed at later time points. This was different to the pattern observed when we used the proteasome inhibitors, where higher and lower bands both responded (although with different sensitivities). Our studies found that phosphorylation of Ube2J1 at S184 is a target for protein phosphatases, with supporting evidence derived from a combination of pharmacological and genetic overexpression studies. Regarding the specific identification of PP2A, it is known to function in the UPR where it has been described to have a role in both mammals (28) and plants (29). Its’ involvement in regulating a translocon component is of particular interest however considering that S184 phosphorylation of Ube2J1 is reported to be of such central importance for UPR recovery (15). In this respect therefore, although our results could simply reflect cellular recovery and a return to homeostasis, they could also indicate a more substantive role in turning off the response. Such a function has previously been proposed for PP2Ce phosphatase, which acts to regulate phosphorylation status of the IRE1 sensor (30).

While our data may point towards a broader role for phosphatase enzymes in regulating ERAD, when it comes specifically to Ube2J1, they are suggestive of a S184-phosphorylated role that requires time limited removal. Although this can be achieved through the activity of PP2A, similar results can be achieved by targeted degradation. Because of the recognised influence of ubiquitinated lysines on proteasomal degradation, we examined whether cis-lysines regulate steady state expression of Ube2J1, and identified K186 as an important residue. Mutating K186 specifically increased the S184-phosphorylated species, yet the mutant still showed partial sensitivity to proteasomal inhibition—an outcome inconsistent with K186 being the sole site required for ubiquitination and subsequent degradation.

Although replacing the C-terminal tail of Ube2J1 with the tail anchor of cytochrome b5 has been shown to be associated with its ubiquitination (16), and a mass spectrometry study suggests that K186 can indeed be ubiquitinated (31), there remains little direct experimental evidence for polyubiquitination of the wild type protein. It remains possible therefore that a non-ubiquitin modification at K186 influences phosphorylation at S184. For example, Carr et al. demonstrated that methylation of pRB at K810 regulates phosphorylation at neighbouring S807 and S811 (32). Indeed, our observations might not necessarily reflect a lysine-dependent mechanism at all: the K186 mutation may simply alter local accessibility of the S184 site.

For Ube2J1, our study provides an insight into the manner in which levels of the S184-phosphorylated protein can be regulated, and point towards a time limited function associated with a specific phase of cellular stress responses. While this could evidently impact on the degradation of misfolding ERAD substrates, it may also influence ERAD effectors, considering that the protein is reported to also regulate steady state levels of components OS9, Derlin 1, Derlin 2 and EDEM1 (33).

## ABBREVIATIONS

Ubc: ubiquitin conjugating
PP2A: protein phosphatase 2A
UPR: Unfolded Protein Response
ERAD: endoplasmic reticulum associated degradation

## ACKNOWLEDGEMENTS

These studies were supported by grant RP/2006/294 to JVF from the Irish Health Research Board (HRB). SYL was a participant in the HRB Cancer Scholars Programme at University College Cork. We wish to thank T.M.Wac for useful suggestions.

## Notes

### Competing Interest Statement

J.V. Fleming served for a period as consultant to Cygenica Ltd.

